# How to define and optimize axial resolution in light-sheet microscopy: a simulation-based approach

**DOI:** 10.1101/783589

**Authors:** Elena Remacha, Lars Friedrich, Julien Vermot, Florian O. Fahrbach

## Abstract

“How thick is your light sheet?” is a question that has been asked frequently after talks showing impressive renderings of 3D data acquired by a light-sheet microscope. This question is motivated by the fact that most of the time the thickness of the light-sheet is uniquely associated to the axial resolution of the microscope. However, the link between light-sheet thickness and axial resolution has never been systematically assessed and it is still unclear how both are connected. The question is not trivial because commonly employed measures cannot readily be applied or do not lead to easily interpretable results for the many different types of light sheet. Here, by using simulation data we introduce a set of intuitive measures that helps to define the relationship between light sheet thickness and axial resolution. Unexpectedly, our analysis revealed a trade-off between better axial resolution and thinner light-sheet thickness. Our results are surprising because thicker light-sheets that provide lower image contrast have previously not been associated with better axial resolution. We conclude that classical Gaussian illumination beams should be used when image contrast is most important, and more advanced types of illumination represent a way to optimize axial resolution at the expense of image contrast.

## 1. Introduction

Light sheet microscopy [1] has proven to be an enabling technology for a number of research fields including developmental biology [2], cell biology [3], neuroscience [4] and many more [5]. Applications benefit from the low photodamage [6] and high image acquisition speed [7, 8] enabled by efficient use of light and high degree of parallelization.

The purpose of the light sheet is to optically slice the sample to generate images with high contrast of thickly fluorescent samples. The thickness of the slice depends on the thickness of the light sheet. In other words, the light sheet fully determines optical sectioning. Illumination by a thicker light sheet results in the projection of a thicker section of the sample into a single image. If the light sheet is thicker than the depth of focus of the detection lens, the image will contain blurred information from object regions illuminated by the light sheet but out-of-focus of the detection lens. In thickly fluorescent three-dimensional samples, optical sectioning (OS) is required to generate image contrast. Because resolution depends on contrast, i.e. the separation of individual features is only possible for a sufficient drop in signal in between two features, optical sectioning should be seen as one of the fundamental parameters of a light sheet microscope.

The length of the light sheet determines the field of view that can be illuminated evenly – with nearly equal optical sectioning and axial resolution. Due to diffraction the thickness depends on the length of the light sheet: A conventional Gaussian-beam light sheet created at a higher NA is thinner at its waist but the distance over which it remains thin is also shorter. In commonly used microscopes that illuminate and detect light using a single objective resolution is anisotropic. In a light sheet microscope, lateral resolution is determined by the NA of the detection objective alone and axial resolution is additionally influenced by the intensity profile of the light sheet, hence by the illumination objective. A quantitative comparison of the resolution of light sheet microscopes and other fluorescence microscopy techniques can be found in [9].

Many researchers put a lot of effort into improving the properties of the light sheet because of its strong influence on image quality. The static light sheet was replaced by a scanned light sheet [10] and holographic beam shaping was employed to generate Bessel beams [11] to reduce scattering artefacts building on their self-reconstruction properties [12, 13]. Since Bessel beams were also found to be non-diffracting which means that their profile does not change over significant distances, attempts were undertaken to use them to provide higher resolution over larger fields of view, which required additional efforts, such as structured illumination or two-photon excitation [14], confocal detection [15] or application of the STED principle [16] to recuperate image contrast. Other non-diffracting beams like the scanned Airy-beams [17] or the scanned Sectioned Bessel Beam [18] but also static line Bessel sheets [19] and static Airy Sheets [20] were also employed. Coherent super-position of Bessel beams, termed lattice light sheet was reported to be especially beneficial [3] by providing high spatiotemporal resolution. In summary, many different light sheet types have been proposed promising benefits especially in terms of achievable image quality.

Considering the abundance of illumination schemes a general measure for comparison of the benefits of the methods is desirable. So far, measures used for the comparison are different across publications and range from image contrast [21], or measures of resolution such as the profile through the image of a bead [3, 9] or the modulation transfer function (MTF) [3, 17, 18]. The lack of a uniform measure or application of measures used in other types of microscopes is a problem for two reasons: Firstly, the properties of the light sheets are very different, and some measures cannot account for these differences. For example, the intensity profile of some light sheets decays monotonically while others show side-lobes. Second, and more importantly, these measures may not be adequate to reflect the influence of parameters that control the ratio of axial resolution and image contrast that they provide. For example, in a confocal microscope, axial resolution and image contrast are linked unambiguously, i.e. they cannot be influenced independently. But in a light sheet microscope, depending on the light sheet employed resolution and contrast can be controlled largely independently. Accounting for this additional degree of freedom, we aim to provide measures to quantitatively analyze and compare light sheets in this publication.

Here, we establish a set of versatile and accurate measures that take the special properties of light sheet design into account and allow to quantify properties that are directly relevant for the image quality: axial resolution and optical sectioning. The measures were applied to a selection of different light sheets. Through this comprehensive assessment we gained understanding of the trade-offs between different illumination approaches. The question to be answered in this paper is not primarily which beam is better than the other because we found a simple binary answer to be inadequate. Rather, this paper aims to provide the reader with measures and a framework applied to a neutral and rigorous comparison between the trade-offs that exist between different illumination concepts in light sheet microscopes to enable an informed decision. This endeavor is supported by the Matlab® code we provide with this publication that can be used to simulate various existing light sheets as well as new ones and quantitatively compare them to each other.

We apply our measures to simulated data of the 3D intensity distribution of the different light sheets for linear and two-photon excitation. The choice in our analysis was made according to the following rules: We limited our analysis to light sheet types that depend on the NA and one additional parameter. The beams differ in the order of the polynomial that describes the phase in the focal plane: A linear function (φ(*r*) ∼ *r*) for the Bessel beam, the Lattice Light sheet and the Double beam (φ(*x*,*y*) ∼ *x*), a quadratic function (φ(*r*) ∼ *r*^2^) for the focused flat top and the Gaussian beam, a cubic function (φ(*x*,*y*) ∼ *x*^3^ + *y*^3^) for the Airy beam, and a quartic function (φ(*r*) ∼ *r*^4^) for the Spherically Aberrated beams. Light-sheets can be generated with these beams either by scanning across the field of view or by shaping the beam directly into a sheet, for example by a slit aperture in the back aperture of the illumination objective. In principle, lattices can be generated by a grid in the back focal plane for any kind of beam, not only the Bessel beam. When formed into a sheet by a slit aperture, the profile of the Double beam, the Focused flat-top beam, the Gaussian beam, the Spherically aberrated beam remains unaffected. We did not include scanned Bessel beams because we already included two light-sheets with linear phase and Bessel beams were already thoroughly compared and found to be inferior to the Lattice Light Sheet [3, 25].

Our simulation is based on the scalar propagator approach [22] which considers monochromatic, coherent light. The propagator itself represents an analytical solution to the Helmholtz-equation and does not use approximations. We validated the accuracy of the simulation, especially to ensure that apodization and polarization effects can be neglected for light sheets [23, 24]. Details on the simulation are explained in Appendix B.

## 2. Characterization of light sheets

In this section we give a short introduction to illumination beams that have been employed for sample illumination in light sheet microscopes. An overview of the different light sheets is shown in Figure 1. The beams are described in more detail in Appendix A. Conceptually, a division into two major categories is helpful:

**Figure 1:**
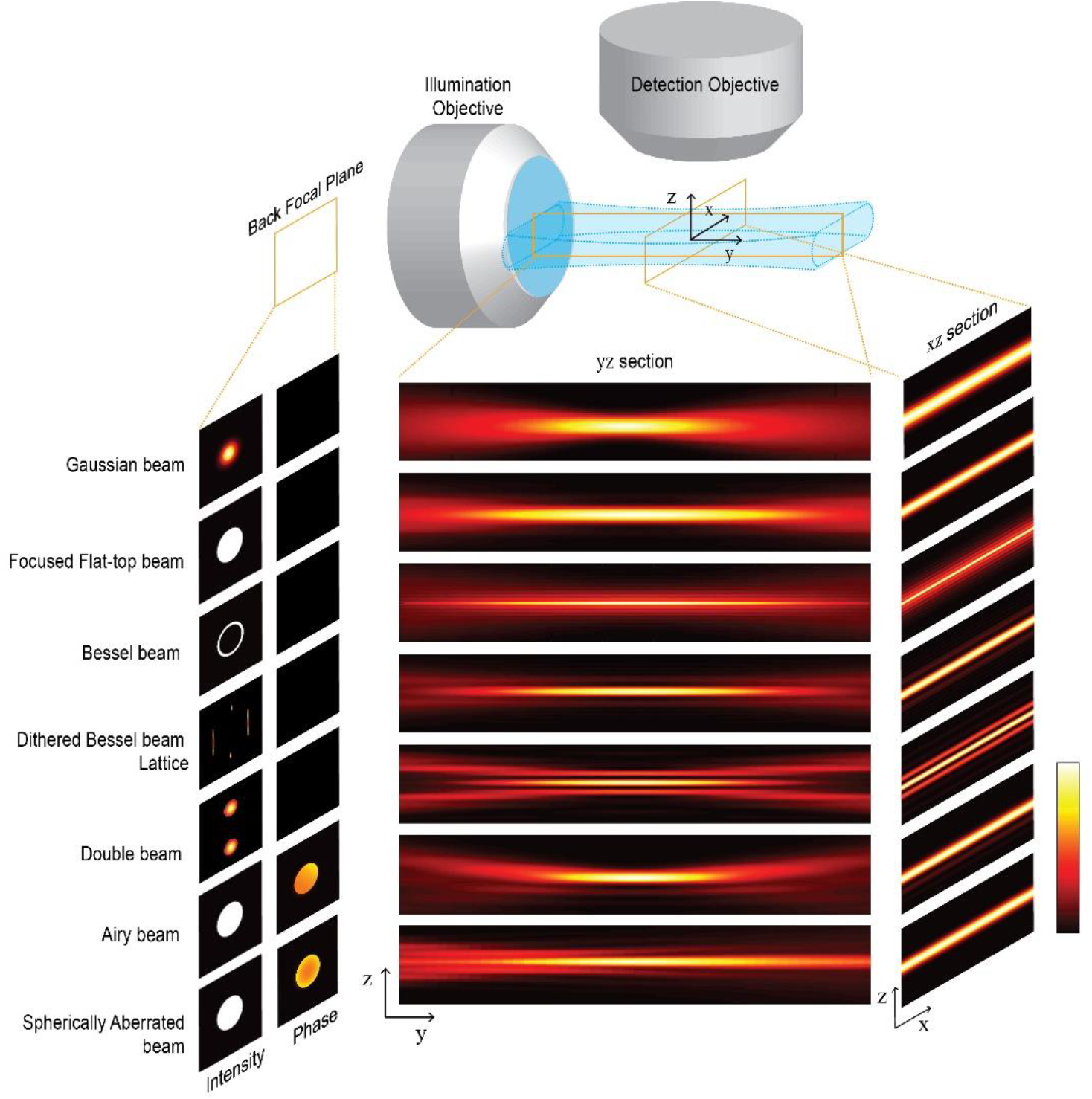
Illustration of the six light sheet types included in the comparison. The amplitude and phase of the electrical field in the back focal plane of the illumination objective is used to simulate the intensity distribution of the beams in the sample volume. Cross section along the illumination and detection axis (YZ) and along the scan axis and detection (XZ) of the sheets resulting from beam scanning or sheet dithering.

- Single-lobe beams: These beams can be described by a single parameter: the focusing NA. The most commonly used such beam is the Gaussian beam [1]. The intensity profile decays monotonically from the peak following a Gaussian distribution. The profile of Focused flat-top beams is often referred to as Airy disk. It exhibits a strong peak and very weak sidelobes and the ratio of their respective magnitudes is fixed.
- Multi-lobe beams: These beams are determined by two parameters. Examples constitute the Bessel beam, the Dithered Square Lattice Light Sheet [3], Airy Beams, [17, 20], Spherically Aberrated beams and the Double Beam. Light-sheets can be formed statically, e.g. by a cylindrical lens in the beam path, by dithering the lattice or scanning the beam in the focal plane of the detection lens. The intensity profile of multi-lobe beams does not decay monotonically with the distance from the main lobe but shows a socket (e.g. for Spherically aberrated Beams) or exhibits side lobes or side sheets (e.g. for the other three types). The auxiliary structure carries a large proportion of the total power of these beams.

## 3. Optical performance parameters of light sheet microscopes

Here, we introduce three measures for the optical performance of light-sheet microscopes: Two for thickness and one for length. Most importantly, we use two separate measures to quantify different aspects related to the thickness of the light sheet that have different meanings for the performance of the microscope. This separation is both necessary and suitable to characterize a wide range of light sheets.

### 3.1 Quantitative Analysis of light sheets

In this section we explain the measures we establish in this manuscript and explain their practical meaning and relevance to the performance of the microscope.

#### 3.1.1 How thick is a light sheet? Main Lobe Thickness vs Optical Sectioning

When assessing the thickness of the light sheet it is important to keep in mind the two image quality parameters that the light sheet affects: axial resolution and image contrast. We base our analysis on the facts that:

- axial resolution depends on the width of the profile of the microscope’s point-spread function (PSF) along the detection axis, and
- image contrast depends on the overall thickness of the light sheet.

Both parameters should be reflected by measures to be able to compare the performance of different light sheets. A measure for axial resolution should be based on the width of the light sheet’s intensity profile along the detection axis. Contrary to this, the second measure for the thickness of the light sheet should quantify the range along the detection axis over which a significant portion of the total detected fluorescence originates.

For a conventional light sheet created by a Gaussian beam the two parameters are closely related and can be assessed with one existing measure. For example, the waist of a Gaussian beam is defined as the range around the peak where the intensity is larger than 1/e^2=13.5% of its value at the peak. The beam carries 1-1/e^2=86.5% of the power within a circle of that radius. Both measures are inseparably linked. For other beams, the measure needs to be generalized to reflect both the drop in intensity around the peak and the region carrying a substantial fraction of the power.

Looking at a light sheet formed by a scanned beam the relevant quantities are

- the distance along the detection axis where intensity drops to a fraction of the peak intensity and
- the total fluorescence generated by the light sheet within a certain distance from the detection focal plane.

The first measure yields axial resolution while the second yields contrast. Figure 2a shows an example for a light sheet formed by a single lobe beam. The width of the profile measured at 1/e=37% of the peak intensity is shown in Fig 2c, d for two positions along the beams propagation axis and the integral under the profile that marks 1-1/e=63% of the total power of the beam in shown in Fig. 2g, h. Within the range where the intensity is larger than 37% of the peak intensity, the light sheet with a Gaussian intensity profile carries a higher fraction of 84% of its total power. The reason is that the light sheet is not a circularly symmetric beam but essentially spreads in only one direction. For the purpose of comparing different light sheets the exact ratio is not essential, but the fact that this ratio cannot be influenced. Most importantly the ratio is independent of the length of the light sheet. The first (drop in intensity) and second measure (power contained within a region) lead to the proportional results for any length of the single-lobe light sheet.

**Figure 2:**
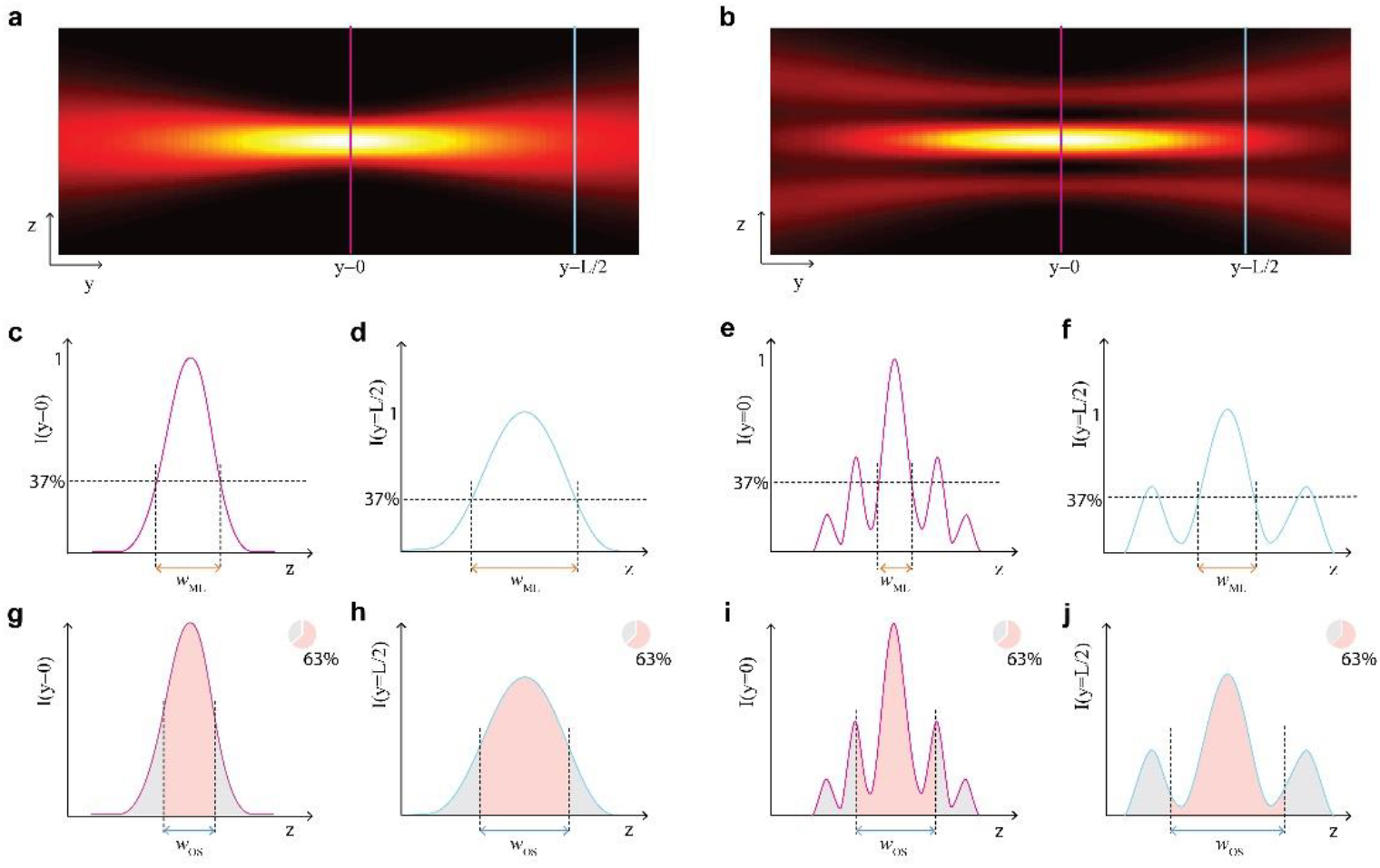
Definition of Main Lobe Thickness and Optical Sectioning. *yz*-cross sections of the intensity of a single-lobe beam (a) and a multi-lobe beam (b) are shown. The respective profiles at the focus (purple line) and at a distance *y*=*L*, where the thickness has doubled (blue line) are shown in c-j. c, d and e, f illustrate the main-lobe thickness, defined as the range where the intensity is higher than 37% of the maximum intensity in the focus, in focus (c, e) and at distance *y*=L (d, f) for single-lobe beam (c, d) and multi-lobe beam, respectively. Sub-figures g-j correspondingly illustrate Optical Sectioning *w*_OS_, defined as the range that contains 63% of the beam’s power.

However, for multi-lobe light sheets that exhibit a more complex structure the values obtained by the first and the second measure may be substantially different. As indicated in Figure 2 on the right side, measuring the distance over which the intensity profile drops to 1/e of its peak captures only the thickness of the main lobe (Fig. 2e, f). Therefore, we introduce the term “main lobe thickness” for this measure. Optical Sectioning *w*_OS_, indicate in Figure 2i,j considers that the main lobe may not carry sufficient power to contribute 63% of the total detected signal. This measure gives a substantially larger value than the main-lobe thickness and more closely reflects the true light sheet thickness.

In many previous publications a missing distinction between light sheet thickness and thickness of the main lobe could cause confusion, e.g. when referring to „ultrathin planar illumination produced by scanned Bessel beams” or “Ultrathin Bessel Light Sheets” when actually only the main lobe is thinner than a conventional Gaussian light-sheet of equal length [19, 25]. For the most frequently used light sheet with a Gaussian profile there is only one sheet and the measure of main lobe thickness and light sheet thickness are strictly proportional. But as illustrated in Figure 3 this is not true for multilobe beams, that exhibit an auxiliary structure, such as multiple parallel sheets. When only the thickness of the main lobe is considered for multilobe beams a significant fraction of the total illumination power the sample is exposed to is left of consideration. This power contributes a to detected fluorescence signal in a light sheet microscope potentially decreasing the image contrast. It is very important to take into account that while slices through the PSF along the detection axis, that are usually shown, can be used to indicate axial resolution, but they make it very hard for the observer to infer the optical sectioning performance or light-sheet thickness.

**Figure 3:**
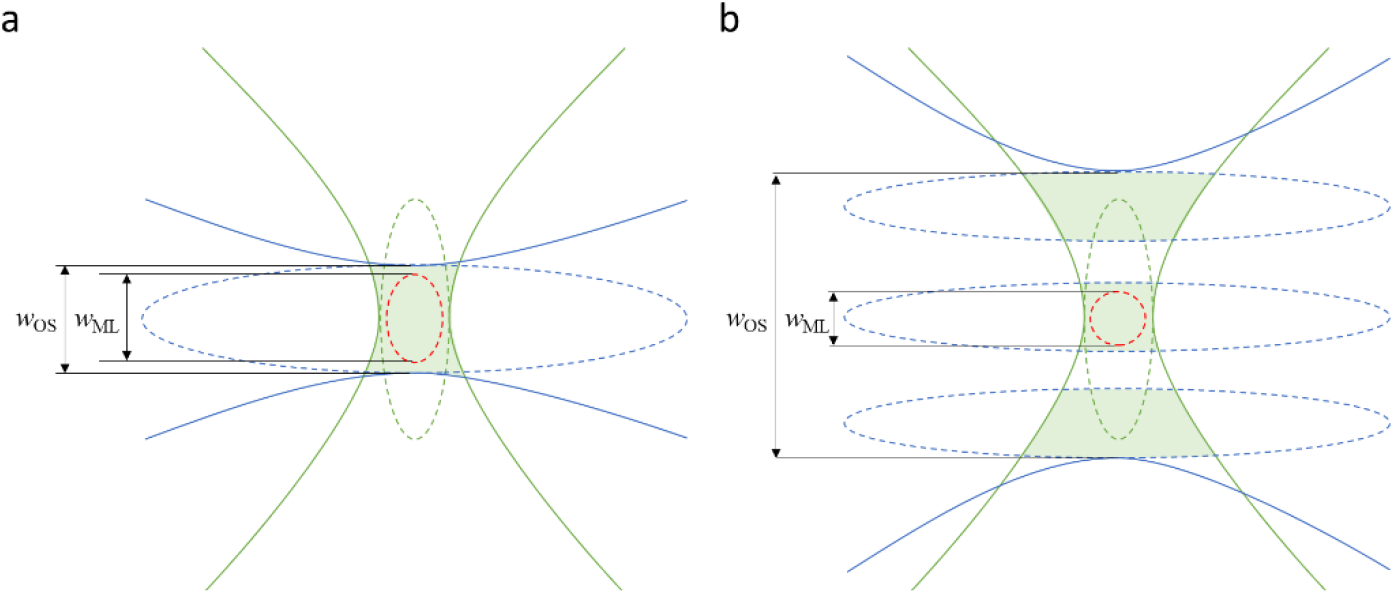
Schematic illustration of the effect of different beam shapes on Axial Resolution *dz* and Optical Sectioning OS in a light sheet microscope. The illumination PSF (blue) indicates the intensity of the illumination light. The detection PSF (green) indicates the location-dependent detection probability for fluorescence photons. The dotted lines indicate iso-intensity surfaces, e.g. at 37% of the peak value. The dotted red line indicates the iso-surface of the combined illumination and detection PSF (product of illumination intensity and detection probability). Its extent along the detection axis is proportional to the axial resolution of the system. The green shaded area indicates the volume where fluorophores are illuminated, and fluorescence is collected. The optical sectioning measures the extent of this volume along the detection axis. In a), where a single lobe illumination beam is shown, resolution is anistropic, i.e. lateral resolution is better than axial resolution, but optical sectioning is close to axial resolution. In b), where a multi-lobe illumination beam is sketched, resolution is isotropic, but additional fluorescence signal is collected from the areas illuminated by the side lobes. Optical sectioning is inferior to axial resolution. The side lobes illuminate out-of-focus planes that blur the resulting image.

#### 3.1.2 How long is a light sheet? Generalized Rayleigh range

Having established a measure for the light sheet thickness, it is sensible to base the definition of the light sheet length on it. This corresponds to a generalization of the concept of the Rayleigh range so that it can be applied to all the beam types included in our analysis. We choose to define the length of a light sheet as the distance over which the light sheet thickness, which we measure in terms of optical sectioning, doubles. This measure yields values equivalent to the Rayleigh range for Gaussian beams (except the Rayleigh range is based on an increase by a factor of √2 and not 2) but better reflects loss in contrast by the spreading of beams with a non-monotonically decaying profile, a broader base or side-lobes. The measure quantifies the minimum extent along the detection axis that contains 67% of the light sheet’s energy regardless of the position of the light sheet’s peak along the detection axis. Even though it would be conceptually easier and more straight-forward to calculate, a measure based on the extent of the on-axis intensity is not useful for the comparison of different light sheets. For single lobe beams, both measures are strictly proportional to each other, but not for the other beams used in light sheet microscopy. In thickly fluorescent samples image contrast is essential and the optical sectioning is the parameter that best reflects the microscope’s ability to generate images with high contrast.

### 3.2 Comparative Analysis

We used numerical simulation to generate the seven different light sheets shown in Figure 1 for both linear and two-photon excitation. For linear excitation we used a wavelength of 488nm and 920nm for two-photon excitation. For each type we varied the length and measured main lobe thickness and full light sheet thickness which is equal to optical sectioning as illustrated in Figure 3. The results are shown in Figure 4.

**Figure 4:**
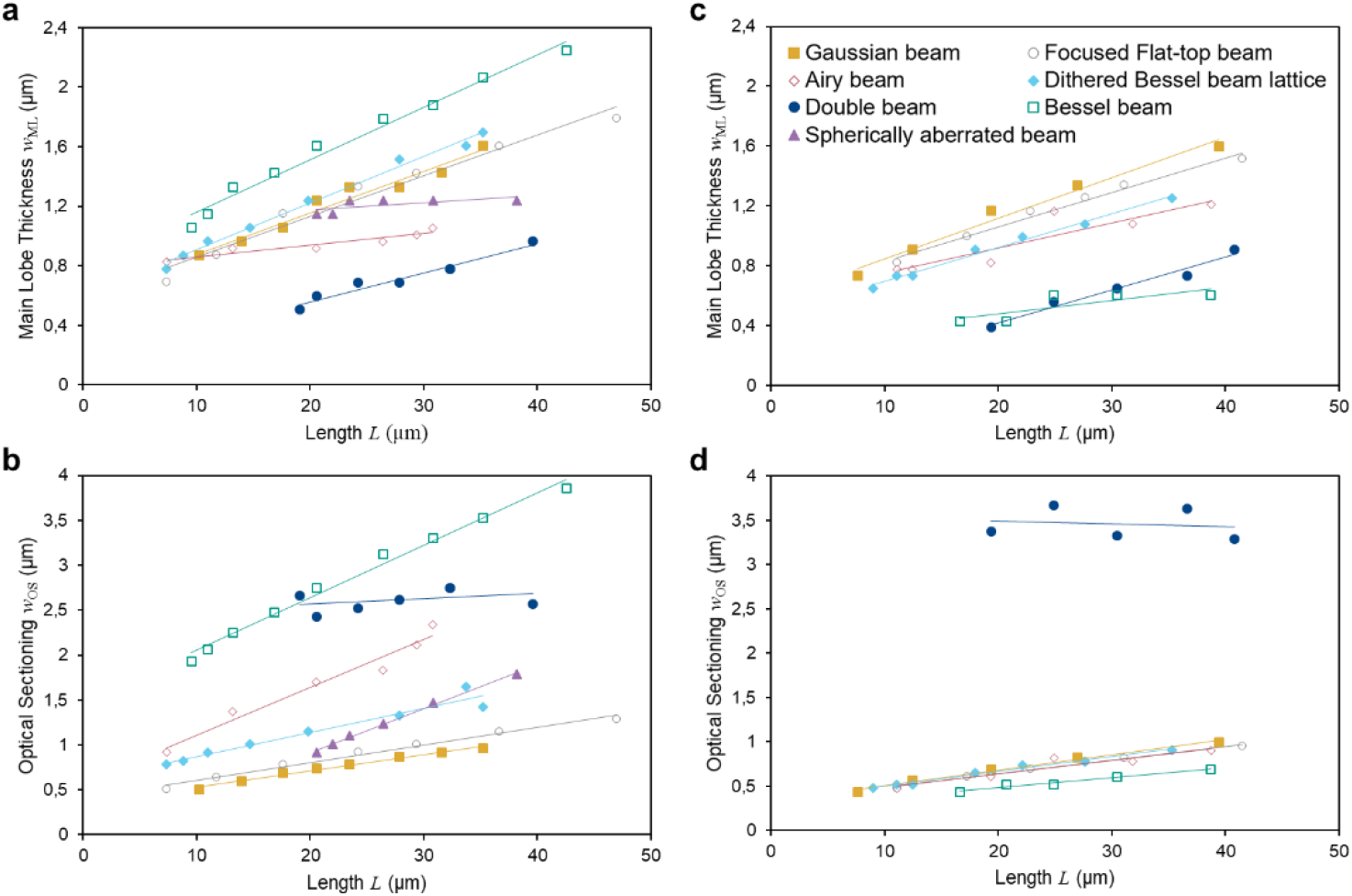
Measurements of main lobe thickness and optical sectioning from simulated data. Main lobe thickness (a, c) and optical sectioning (c, d) are shown as a function of length for linear (a, b) and two-photon excitation (c, d). Parameters used in the simulations are NA = 0.14 to 0.26 for Gaussian beams; NA = 0.16 to 0.40 Focused flat-top beams; NA = 0.05 and separation NA = 0.14 to 0.3 for Double beams; NA = 0.5 and phase parameter a = 4 to 12 radians for Airy beams; NA = 0.32 to 0.6 with ring thickness parameter ε = 0.7 for the Bessel beam, NA = 0.25 with quartic phase amplitude a = 0.04 to 0.09 radians for the spherically aberrated beam; and NA = 0.3 to 0.55 for ε = 0.7 for the Bessel beam lattice.

To tune the length *L* of the light sheet, we varied the Numerical Aperture NA for the Gaussian and Focused flat-top beam. For the Double Beam we adapted the separation between the two sub-beams in the back focal plane, NA_sep_ and held the NA of the sub-beams constant. For the Airy beam, the amplitude of the cubic phase was altered for a constant NA of the beam. For the Spherically Aberrated beam, we varied the magnitude of the spherical aberrations at a constant NA. For the Bessel beam and the Bessel Lattice we varied the NA for constant relative ring thickness parameter. Details on the generation and the beam parameters are given in Appendix B.

We find the expected square-root dependency of both the light sheet thickness and the main lobe thickness on the length for the Gaussian beam and the Focused flat-top beam. For these beams the width depends linearly on the NA and length scales inversely with the square of the NA.

Surprisingly, both the thickness of the main lobe and the optical sectioning of the dithered square lattice light sheet, which was previously identified to provide best confinement of the excitation [3] were found to be a little worse than for the conventional beams. This can be understood when looking at the angular spectrum which consists of two components – the two side bands and a split central band (see Figure 1 where the intensity in the back focal plane is shown). The side bands and the central bands each generate beams that propagate at equal angles with respect to the optical axis. They also have equal depth of field which can be derived directly from the fact that the propagation invariant depth (the depth of focus or length of the beam) is given by the radial width of the angular spectrum [26]. However, the side bands provide less confinement along the detection axis which is a direct consequence of their angular spectra being more confined along the detection axis. The side-bands provide lateral structuring of the light sheet which is useful when structured illumination is used to recuperate image contrast and resolution in a post-processing step, but if the structure is blurred by dithering the light sheet the lateral structure of the light sheet blurs and the sidebands only increase the overall thickness of the light sheet. Another way to understand the dithered lattice light sheet is based on the Field Synthesis Theorem [27]. The dithered light-sheet can also be understood as an incoherent superposition of three light sheets: a line Bessel sheet [19] and two Focused flat-top light sheets with poor confinement along the detection axis.

The spherically aberrated (SA) light sheet represents an interesting example that has not yet been demonstrated in practice. The increase in depth of focus (light sheet length *L*) is obtained for equal NA by increasing the magnitude of the aberrations. At *L*∼15µm the SA is weak and both *w*_OS_ and *w*_ML_ of the SA beam are similar to the diffraction-limited Gaussian beam. However, for stronger SA the length increases and while the increase in main lobe thickness *w*_ML_ is smaller than for the Gaussian beam, the light-sheet thickness *w*_OS_ gets larger. A thicker light-sheet is the price to pay to maintain a thinner main lobe over a larger distance.

Similarly, for the Airy beam *w*_OS_ increases more strongly than for the Gaussian beam while *w*_ML_ increases less. However, *w*_OS_ showed a linear dependency on length *L*. We account this fact to the asymmetrical shape of the beam where all side lobes lie on one side of the main lobe.

The Double beam shows almost no dependency of optical sectioning on length. This effect can be explained by the fact that the Double beam light sheet is the product of the interference of two beams (‘sub-beams’) and the length *L* is, in this case, increased by reducing the angle between the two sub-beams thereby stretching the range along the optical axis where the sub-beams interfere. However, because the NA of the sub-beams are not changed, the cross-section of the interference pattern changes only weakly.

For two-photon excitation we obtained some interesting results. Generally, we expected that the nonlinear fluorescence excitation leads to a suppression of the side lobes and leads to better optical sectioning. The Gaussian and Focused flat-top yield very similar results for linear and two-photon excitation. Here, the longer wavelength is almost completely compensated by the nonlinear response. Interesting results were obtained for the Bessel beam, which performs poorly both in terms of main lobe thickness and optical section for linear excitation but for two-photon excitation performs best of all beams in our selection. In contrast, for the Double beam optical sectioning is even worse for two-photon excitation because the main lobe and the side lobes have very similar intensity. In comparison to Gaussian beams, Airy beams feature better optical sectioning and slightly thinner main lobes.

For all types of light-sheet the optical sectioning and axial resolution they provide depends on their length. Therefore, it is essential to compare light sheet properties only for equal length. Setting the Gaussian beam as the reference value, we compute the relative gain or loss in thickness and optical sectioning for all the different beam types below for the same beam length. Figure 5 shows the results for a depth of field of the illumination beams of approximately 30µm. We conclude that for all beam types there is a trade-off between light sheet thickness and main lobe width. Making a thin light sheet comes at the expense of either length for Gaussian beams or at the expense of optical sectioning for other light sheet types.

**Figure 5:**
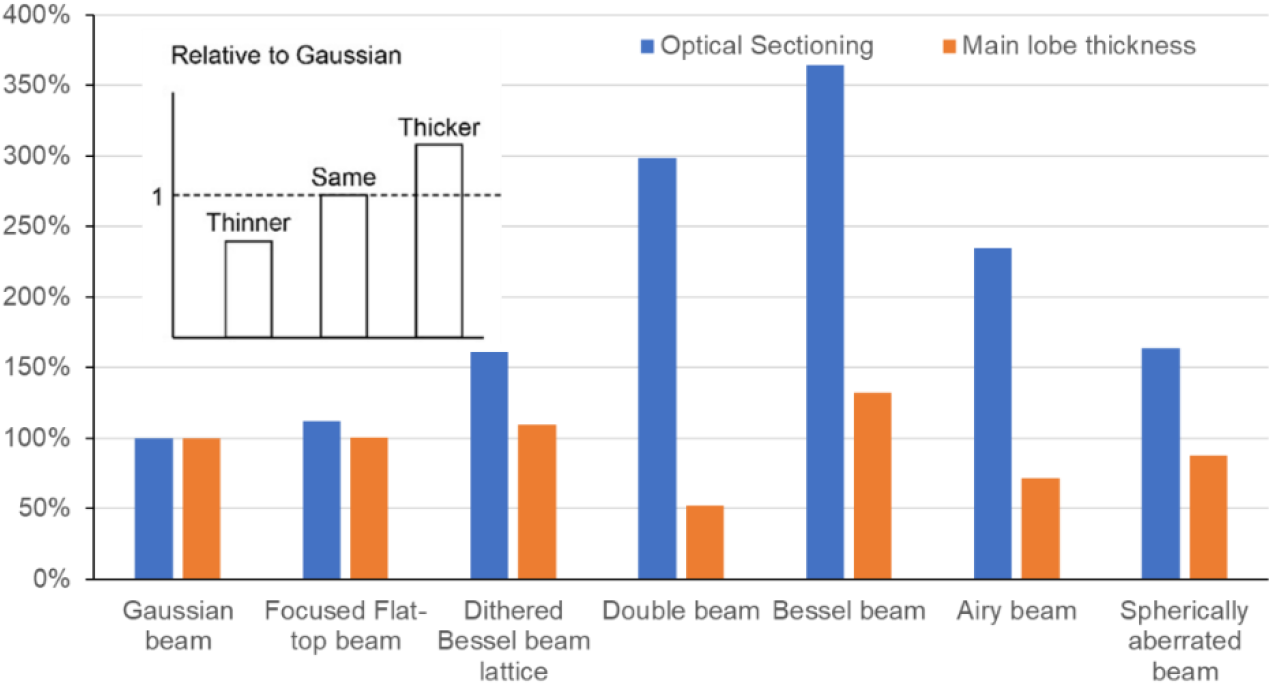
Ratio of Optical Sectioning (blue) and Main Lobe Thickness (orange) for each beam type with respect to the Gaussian beam. All values are computed for beams with length *L*∼30µm. The insert shows a scheme to interpret the graph. Values larger than 100% mean that the main lobe or optical section is thicker than for the Gaussian beam.

In most cases length is the parameter determined by the sample or the field of view that the researcher aims to observe in a single frame. Therefore, we chose the approach to set a light sheet length first and then compare the light sheets according to the axial resolution and optical sectioning they provide and not vice-versa.

To further illustrate the possibility of tuning *w*_OS_ and *w*_ML_ for the multi lobe beams we analyzed the Double beam. The length of the double beam light-sheet can be tuned either by the angle between the two beams or by the thickness of the beams. When the angle is varied, the overall width of the interference region changes only very little because the cross-section of the sub-beams in the plane perpendicular to the optical axis is approximately constant for small angles between the optical axis and the propagation of the sub-beam whereas the range along which the sub-beams intersect along the optical axis changes. However, the smaller angle between the beams results in a larger period of the interference pattern. This effect can be observed in Figure 6 a. It is possible to obtain opposite trends, i.e. almost constant main lobe width and increasing light sheet thickness, if the NA of the interfering beams was altered instead of their interference angle (Fig. 6b). Note that while we used the double beam similar tuning is possible for the Dithered Lattice, the Spherically Aberrated Beam and the Airy beam likewise because the depend on two degrees of freedom (Appendix A & B).

**Figure 6:**
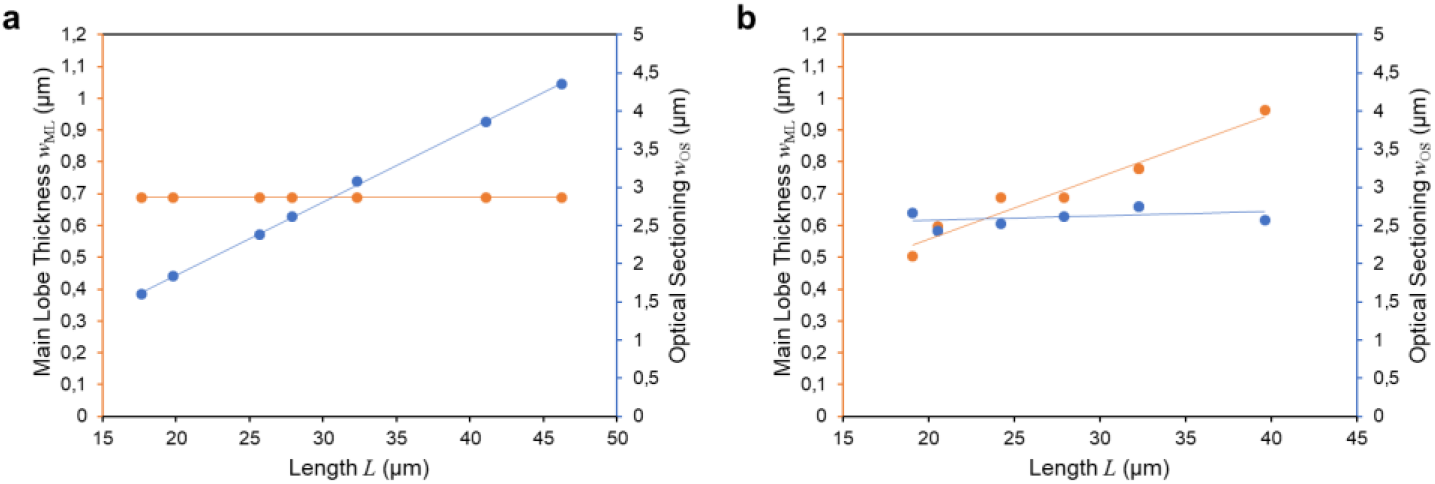
The Main Lobe Thickness (orange) and Light Sheet Thickness / Optical Sectioning (blue) is shown for the double beam. The figure illustrates how the two parameters of multi lobe beams can be used to control beam length at the expense of a thicker light sheet or a thicker main lobe (a) To change the length of the light-sheet the NA of the two sub-beams is varied (NA=0.03 to NA=0.08) while the separation is held constant (NA_sep_=0.2) (b) the NA of the sub beams is held constant (NA=0.05) while the separation is varied (NA_sep_=0.14 to NA_sep_=0.3). A beam with L = 40µm can be reached either by a NA=0.035 and NA_sep_=0.2 or a combination of NA=0.05 and NA_sep_=0.14.

## 4. Discussion

We analyzed the effect of different light sheets on image quality by using three parameters to represent properties which act as the major contributors: the thickness of the main lobe, the overall thickness of the light sheet and the length. Fundamental to our comparison of different light sheets is the insight that it must be performed for light sheets that have equal length. Based on the insight that most advanced light sheets like the Bessel Beam Lattice or the Airy Beam feature two parameters that influence the shape of the light sheet we also implemented two measures: thickness of the main lobe which contributes to the axial resolution and the overall thickness which determines the optical sectioning.

In our analysis we neglected the effect of the detection objective, i.e. its point-spread function. The reasoning for this approach was the following: The detection NA by itself does not affect optical sectioning: Regardless of detection NA the integral collection efficiency of the signal out of any plane along the optical axis of the detection objective is the same. Optical sectioning therefore only arises from light sheet thickness. To get an idea of the influence of the light sheet on axial resolution it is insightful to approximate both the main lobe thickness of the light sheet and the axial profile of the detection PSF by Gaussian profiles: the detection profile is given by

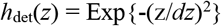

and the main lobe of the illumination profile by

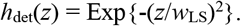

The profile of the resulting point-spread function of the microscope is then

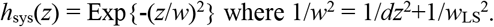

The light sheet always improves axial resolution in comparison to epi-fluorescence illumination. But there is only a significant effect on resolution when the light sheet’s main lobe thickness is smaller than the depth of focus of the detection lens. Some light sheet with thin main lobes may therefore provide higher axial resolution: For example, for equal depth of field and light sheet thickness, the resolution is increased by 30% (for *w*_LS_ = *dz*: *w* = 0.7 *dz*) but only by 10% for a light sheet that is twice as thick than the detection objectives depth of field (for *w*_LS_ = 2 *dz*: *w* = 0.90 *dz*), something that routinely happens for detection NA_det_=1.0 and a light sheet created at NA_ill_=0.1 which results in a depth of focus of *dy*_LS_ = 2*nλ*/NA^2^ ≈130µm. However, one important parameter that depends on the detection PSF is image contrast: The relationship between the optical section provided by the light-sheet and the depth of focus of the detection objective determines image contrast (see Figure 3). If the light sheet thickness is larger than the detection depth of focus out-of-focus planes within the sample are illuminated and detection and image quality is reduced. In conclusion, taking the detection NA is not required for a comprehensive comparison between different light sheets even though it influences image quality.

Image post-processing plays an increasingly important role in microscopy. Deconvolution represents a powerful mean to computationally reassign the detected photons to their original position and thereby greatly increase image contrast in images blurred by out-of-focus signal. Any light-sheet where the optical sectioning is larger than the main lobe thickness will benefit from post-processing. Deconvolution is especially helpful when the depth of focus of the detection lens is shorter than the optical sectioning some of the illuminated structure will be out-of-focus and blur the image. For the selection of light-sheets the option of post-processing means the following: When the quality of raw data matters, beams with good optical sectioning are preferable. When deconvolution is employed, it may be preferable to use light-sheets with thinner main lobes even if they have inferior optical sectioning because the high resolution provided by the main lobe can be recuperated. However, there are limitations to the amount of background that deconvolution can handle. For example, the signal generated by the main lobe should not be covered in the noise of the background signal generated side lobes of the light sheet. In practice, for example a Double beam light sheet can be generated to be both very long and have a thin main lobe, but the strong background from poor optical sectioning will make deconvolution impossible. Moreover, the additional photodamage from thicker overall light-sheets has also to be considered. Our method offers guidance for the choice of ideal imaging conditions: which parameters can be used to generate light sheets for a given sample size that provide images with ideal axial resolution without too much background for deconvolution.

The analysis of the light sheet could also be based on the MTF or OTF as in previous publications [3, 18]. A similar approach, but based on image data, has been used previously [21] but its use to quantify the axial resolution is not straightforward. To the trained eye, the MTF reveals at first glance the maximum spatial frequency representing smallest details that can be resolved. The contrast that these details will hold in the image is visually less easy to quantify because it is proportional to the quotient of the area under the MTF curve at high-frequencies to the area at low frequencies. The MTF is a well-established standard measure for the optical quality of a system to those comfortable thinking in k-space. Our set of measures is meant to provide a complementary approach in real space. We expect it to be a more intuitive method to quantify and compare the optical performance of light-sheets, especially for the many users of microscopy with a background in biology and medicine rather than optics.

We used simulation data to compare the beams. Since new beams are continuously introduced in the field we see the benefit that by the simulator we provide a tool that can be easily used to assess performance of emerging light sheets with existing beams. The simulation method itself has been validated several times e.g. by [23, 24]. It is reliable and accurate. An alternative simulation method that uses Debye theory is presented in [28]. The beam propagation method (BPM) represents an extension to the propagator approach which uses approximations but works in optically inhomogeneous media [12]. The BPM has been used in a software package termed BioBeam that simulates light propagation through optically inhomogeneous media both in the illumination and detection path for different light sheet types at greatly improved speed by relying on a GPU [29]. In principle, our analysis can be readily applied to BioBeam results as well to assess performance in inhomogeneous media.

We showed that two-photon excitation generates much better optical sectioning without compromising axial resolution for many light-sheets, especially the Bessel beam but not the Double beam. However, even though the side lobes do not excite fluorescence they still carry a substantial fraction of the total power of the light sheet. The sample is therefore subjected to a much larger total power than for Gaussian beams.

Other, more complex, illumination and detection schemes are suitable improve axial resolution and optical sectioning at the same time. Examples include structured illumination [14], confocal slit detection [15], STED, RESOLFT [30], tiling [31], axially scanned light sheets [32] and computational reconstruction of images acquired with complementary light sheets [33, 34]. However, the improved performance comes at a cost such as, for example, excess light exposure or reduction in achievable imaging speed. Image quality is controlled by additional parameters and degrees of freedom like the slit width for confocal slit detection or axially scanned light sheets or the power of the de-excitation beam for STED and would therefore require the extension of our model. We limited the selection of light-sheets to those that depend on only two parameters and see an extension of our method to more parameters as a very interesting direction for further development, potentially building on the Matlab® code that we provide which is simple, easy to use and enables interested readers to simulate and analyze not only the light sheets used in this publication but also arbitrary ones.

## 5. Conclusion

In conclusion, we highlight the difference between the optical sectioning and the axial resolution that different light sheets provide. To this end, we distinguish between main lobe thickness and light sheet thickness resulting from the main lobe and the auxiliary structure when comparing different light sheets. We assessed the optical performance for seven different light sheet types by quantifying three different parameters, main lobe thickness, optical sectioning and length. Surprisingly we found no clear winner: none of the light-sheets we analyzed is suitable to provide improved axial resolution and optical sectioning at the same time. Multi-lobe beams have two parameters that can be used to trade optical sectioning, i.e. image contrast against axial resolution. Our publication is meant to offer guidance which choice of beam and parameters may be best for a certain application depending on whether axial resolution provided by a thin main lobe of the light sheet or image contrast and efficient use of light provided by thin optical sectioning is more important, for example in localization microscopy [35]. Axial resolution can also be recovered from thick light-sheets by image processing such as deconvolution. However, the sheet must exhibit a thin main lobe, the signal from that main lobe should stand out from the noise of the background generated by the auxiliary structure of the sheet and the sample needs to tolerate the additional light-dose from the auxiliary structure. Based on the results from our analysis we suggest using Gaussian illumination beams for large FOVs, light-sensitive or thickly labelled samples where efficient use of light is key and therefore best optical sectioning is most important. For high flexibility and resolution in sparsely labelled samples we recommend the Double beam which we found to offer the best trade-of between optical sectioning and axial resolution that can be generated efficiently and where the trade-off between main-lobe thickness and light sheet thickness can be easily controlled.

## Acknowledgements

The authors acknowledge helpful discussions with Nicola Maghelli, James Manton, Petra Haas and Werner Knebel.

## Appendix A: Details on the light sheets

### Gaussian beam

Laser beams ideally emit Gaussian beams. Gaussian beams are characterized by having an intensity profile along the radial direction following a Gaussian distribution. This profile is conserved along the propagation direction, only changing the beam radius. Light sheets with a Gaussian profile can be generated by a cylindrical lens [1] or scanning a Gaussian beam in focal plane of the detection objective [10]. This type of light sheet is by far the most common.

### Focused flat-top beam

A Focused flat-top beam is generated by homogeneously illuminating the back focal plane of an objective lens. The resulting intensity distribution in the focal plane is termed “Airy Disk” which is not to be confounded with the Airy beam and characterized by a main lobe with a very weak ring structure. Contrary to Gaussian beams, the shape of the intensity profile is not conserved along propagation. In practice a Focused flat-top beam is generated by expanding a Gaussian beam by a large factor so that its diameter is much larger than that of the back focal plane (BFP) of the objective lens. The profile can be therefore be assumed to be approximately even across the BFP. For a given NA of the objective lens it is sensible to crop the Gaussian beam by the aperture of the objective so that a larger fraction of the light is sent into the focus at a large angle with respect to the optical axis thereby leading to a tighter focus, at the expense of an additional ring structure. Focused flat-top beams can be used in confocal microscopy to increase lateral resolution because the fluorescence excited by the ring structure is suppressed on the detection side by the pinhole.

### Dithered Lattice

The Lattice light sheets is a coherent superposition of equidistantly spaced Bessel beams. Depending on the distance at which the beams are separated different interference patterns are created. The main two are called square lattice and hexagonal lattice, names that stem from the light distribution in the back focal plane. Here we will consider only square lattices, which are recommended to be used in the dithered mode.

### Airy beam

Airy beams have a very characteristic banana shape. Experimentally, they can be generated by imposing a cubic phase to a Gaussian beam. This can be done, for example using a spatial light modulator. They are self-reconstructing, which means that if they find obstacles in the way they are able to recover their initial intensity profile. They were introduced to light-sheet microscopy for biological research by [17]. They can be used in two different orientations with different advantages and disadvantages. In one configuration, the beam’s main lobe is curved within the detection focal plane, but the beam’s side lobes lie on either side of the main lobe. The advantage is that the main lobe illuminates a flat plane and the disadvantage is that the side lobes on either side of the main lobe lead to a thicker overall light sheet. If the beam is rotated by 45° around the propagation axis, the main lobe of the sheet is bent with respect to the detection objective’s focal plane, but some lobes are situated next to the main lobe along the scan direction, i.e. they illuminate the same volume as the main lobe and thereby do not decrease optical sectioning. We included this version in our analysis because it provides better values in terms of optical sectioning. However, this approach requires heavy postprocessing to correct for the curvature of the illuminated volume. Additionally, a detection lens with sufficient depth of field (low NA leading to low lateral resolution) and/or post-processing by deconvolution need to be considered.

### Spherically Aberrated Beam

Spherical aberrations beams occur when an objective lens designed to focus collimated light is used with defocused or focused light generating away from the objective’s native focal plane. Most commonly, spherical aberrations occur in the presence of refractive index mismatch, i.e. an objective designed to focus into a medium with a refractive index is used to focus into a another medium with a different refractive index. Differential refraction of the rays depending on their radial position translates in an elongation of the beam’s peak and the formation of a socket around it. This type of beams has not been demonstrated for illumination purposes in a light-sheet microscope before, but the concept of focus elongation by spherical aberration has been used to increase the depth of focus of detection objectives in microscopy [36]. We model it by imposing a radially dependent biquadratic phase mask. Note that this beam is the only beam in our selection where the profile along the illumination axis is asymmetrical.

### Double Beam

The Double Beam is an interference pattern created by two beams (“sub-beams”) focused by an objective which then propagate at an angle relative to each other and to the objectives optical axis and interfere in the focal plane of the objective. The beams thereby generate an interference pattern consisting of alternating stripes of maximum and minimum intensity like Cosine Gauss beams [19]. Effectively the double beam generates a stack of light sheets. For light-sheet microscopy the central, and strongest light-sheet is placed in the focal plane of the detection lens. The auxiliary light sheets illuminate parallel planes [37]. The period of the interference pattern is equal to the separation of the light-sheets and depends on the angle of propagation between the beams. The depth of field on the diameter of the beams and the angle. Double Beams can be generated by splitting a laser beam, e.g. a Gaussian beam, and sending the two sub-beams it into the back focal plane symmetrically around the center. The distance between the beams in the back focal plane determines the angle of propagation of the sub-beam and the diameter determines the sub-beams depth of field, i.e. the length of the interference pattern along the objective’s optical axis.

## Appendix B: Details on simulation method

We performed simulations of the propagation of light using the Beam-Propagator Method. The intensity distribution of a light sheet is expressed by the illumination point spread function (PSF) *h*_ill_. In combination with the detection PSF *h*_det_, *h*_ill_ gives the response of an imaging system to an infinitely small point source. The illumination PSF *h*_ill_ can be calculated from the electric field *E*(*x*,*y*,*z*) as:

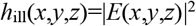

*dy* from the focus as:

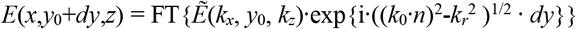

Where *k*_0_=2π/λ is the vacuum wavenumber and 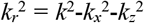 is the radial component of the wave vector *k*. Here we assume that the light is propagating in water and therefore the refractive index *n* is constant and equal to 1.33. An important aspect of the simulation is the choice of discretization. It must be ensured that the sampling *Ẽ* in k-space, *dk*, and *E* in real space, *dx*, are smaller than the relevant features. Because both values are inversely proportional to one another (dx = 2 π / N dk) where *N* is the length of the array, the simulation of beams with either a very narrow spectrum (low NA) or small foci (high NA) requires large arrays. The method works most effectively for medium NAs where both the spectrum and the beam are broad enough to be sampled well without the need for large arrays (large N).

The electric field at position of the focus, *y*_0_, can be in turn computed as the Fourier Transform of the field in the Back Focal Plane (BFP) as:

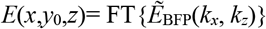

In our simulations, we defined the amplitude and phase of the field in BFP depending on the type of beam used, calculated the electric field at position *y*_0_ and propagated the field. Due to sampling requirements, very long light sheets that require very large arrays were not considered.

Apodization and polarization can also be considered by multiplication *Ẽ*_BFP_ with pupil functions [23, 24]. However, both effects don’t lead to significant changes in the shape of the PSF only for NA>1.0 (n=1.33) and these values are not reached by illumination beams in light-sheet microscopy. Even for the beams with the highest NA shown in this manuscript (focused flat top beam at NA=0.44 and lattice light sheet at NA=0.6) we did not measure changes to the measured quantities by considering the sine condition and performing a vectorial computation. Short pulses used for two-photon excitation have partial coherence which the simulation does not cover. Therefore, aspects depending on pulse the length such as the quantitative relationship between pulse energy and fluorescence excitation cannot be evaluated but do not affect the analysis of different light sheet shapes.

Depending on the number of parameters needed to define the amplitude and phase of the field we can classify the beams in one-parameter or two-parameter beams.

### One-parameter beams

- Gaussian beam: In the BFP, the amplitude follows a gaussian distribution along radial profile. The phase is constant throughout the aperture.

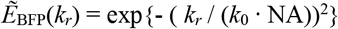
- Focused flat-top beam: Amplitude and phase are constant over the aperture.

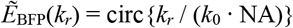

### Two-parameter beams

- Bessel beam The Bessel beam is generated by a ring-shaped intensity. The inner radius is given by √ε *k*_0_ NA and the outer radius by *k*_0_ NA.

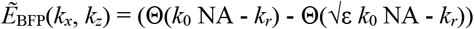
- Lattice light sheet: The lattice is generated by the product of a ring-shaped intensity with a grid structure with period δ*k*_*x*_. The ratio between thickness of the ring and its radius is 1-√ε. The field in the back focal plane is given by:

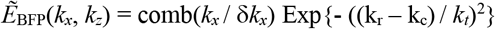 With *k*_*c*_ = *k*_0_ NA (1+√ε) / 2 and *k*_*t*_ = *k*_0_ NA (1-√ε) / 2 Square lattice results from δ*k*_*x*_ = √ε *k*_0_ NA, hexagonal lattice from δ*k*_*x*_ = √ε *k*_0_ NA / 2
- Airy beam: A cubic phase mask is superimposed on a homogeneous illumination in the BFP with

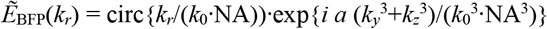

where the amplitude of cubic phase is controlled by *a*.
- Spherically aberrated beam: The amplitude is constant on the aperture. The phase follows a biquadratic function.

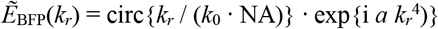

where the exponential term gives the phase modulation, and *a* stands for the amplitude of the phase difference.
- Double beam: The double beam is generated by two gaussian beams determined by NA in the back focal plane centered at positions +δ*k*_*z*_ and -δ*k*_*z*_ where δ*k*_*z*_ = *k*_0_ NA_sep_ with

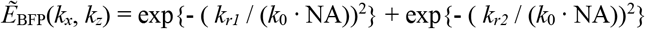

where 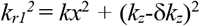 and 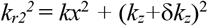.

## Appendix C: Manual to the simulation software

Our light sheet simulator app numerically computes the 3D intensity distribution of light sheets and quantifies its dimensions. It can generate Gaussian beam, Focused Flat Top beam, Airy beam, Bessel beam, Bessel beam lattices, Double Beams and Spherically Aberrated beams, as well as arbitrary types of light sheet introduced by the user. The algorithm takes as initial conditions the amplitude and phase of the electric field of the beam in the back focal plane and calculates its propagation around the focal plane using the beam propagation method.

The program can be downloaded from www.github.com/remachae/beamsimulator. To operate it, run LightSheet_Simulator.mlapp in Matlab environment. There are three possible actions:

- Simulation: Allows visualization of cross-sections of the light sheet as well as the intensity and phase in the back focal plane. To simulate the light sheet discussed in this work, select the beam type in the tab panel and type in the required parameters. To simulate a custom light sheet, select the Create Light sheet tab. Type in the paths to the files containing the amplitude and the phase in the back focal plane. The files must be provided in tiff format.
- Analysis: After simulation of the light sheet, press “Analyze” to obtain the values for Main Lobe Width *w*_ML_, Optical Sectioning *w*_OS_ and Length *L* as defined in this work.
- Saving: Creates the file simulated_ligthsheet.m in the directory indicated under Save workspace. The file contains a 3D intensity matrix and the numerical and physical parameters used in the calculations.

**Figure A:**
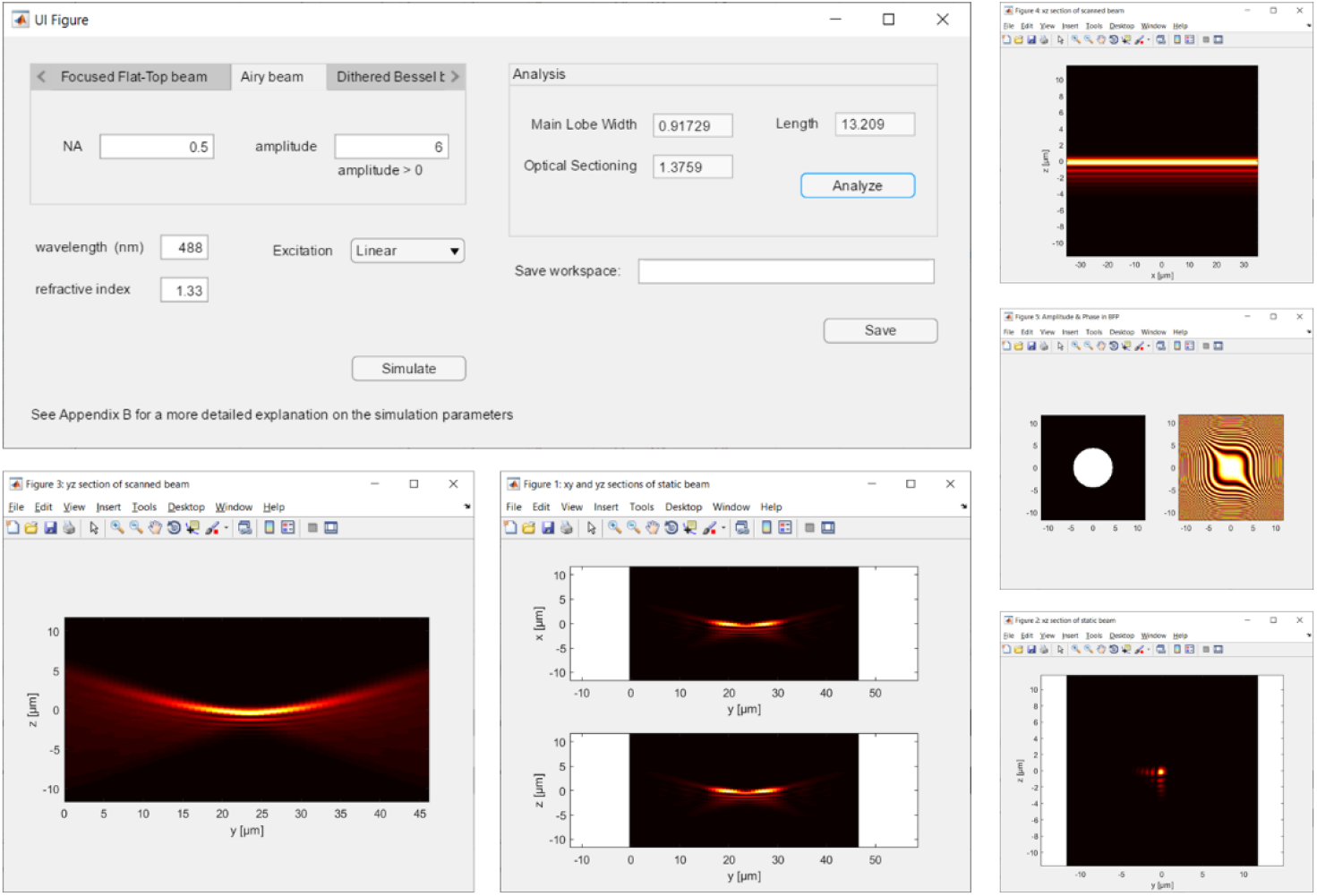
User interface of the beam simulator.

